# Energy-based Advection Modelling Using Bond Graphs

**DOI:** 10.1101/2022.07.07.499159

**Authors:** Peter J. Gawthrop, Michael Pan

**Affiliations:** Systems Biology Laboratory, School of Mathematics and Statistics, and Department of Biomedical Engineering, University of Melbourne, Victoria 3010, Australia; School of Mathematics and Statistics, Faculty of Science, University of Melbourne, Victoria 3010, Australia

**Keywords:** Systems biology, advection, physiome, bond graph

## Abstract

Advection, the *transport* of a substance by the flow of a fluid, is a key process in biological systems. The energy-based bond graph approach to modelling chemical *transformation* within reaction networks is extended to include *transport* and thus advection. The approach is illustrated using a simple model of advection via circulating flow and by a simple pharmacokinetic model of anaesthetic gas uptake.

This extension provides a physically-consistent framework for linking advective flows with the fluxes associated with chemical reactions within the context of physiological systems in general and the human physiome in particular.

## 1 Introduction

Advection is the transport of a substance by the flow of a fluid. Modelling the Physiome [1–3], in particular modelling circulatory systems [4–6], requires methods for modelling the molar flows due to advection. Bond graphs have recently been used in Physiome modelling due to their ability to model a wide range of biophysical systems [5]. To date, the bond-graph approach to modelling biomolecular systems [7–9] considers the *transformation* of molecules via chemical reactions and the corresponding molar flow rates.

However, as pointed out by Cellier [10, p.397] “The term ‘molar flow’ has … been used in two quite different contexts. On the one hand, it denotes the *physical transport* of matter from one point in space and time to another, while on the other hand, it describes the *transformation* of one chemical species into another during a chemical reaction.” In fact, both types of molar flow are needed in physiological models. For example, the binding and unbinding of oxygen dissolved in blood to haemoglobin can be viewed as a chemical reaction leading to transformation molar flow; however, the *advection* of haemoglobin though the blood stream transports the haemoglobin from the lungs to other organs and back again leading to advection molar flow. There is therefore a need to bring both transformation and advection under a common modelling framework; this paper shows how the bond graph approach can provide this framework.

Previously, *Convection Bonds* [11, 12], which carry *two* effort variables, have been proposed and advection has been modelled in the context of chemical reactors by Couenne et al. [13]. In contrast, this paper uses the standard bond graph representation with a single effort variable thus allowing standard bond graph notation and software to be utilised. Although energy-based modelling of distributed (PDE) systems is possible [14], this paper takes the lumped approach which is typically used for modelling the cardio-vascular system [15–17] and allows the use of well-established fast ODE solvers.

§ 2 examines hydrochemical transduction – the transduction of energy between fluid flow and advected chemical potential – from a bond graph perspective. § 3 looks at the advection of chemical species though an orifice and though a pipe and suggests a new bond graph component: **RA**. § 4 compares and contrasts advection of chemical species via the **RA** component and transformation of chemical species via the bond graph **Re** component. § 5 shows how circulatory advection (such as the human blood circulation) combines with binding and unbinding of a ligand and enzyme (such as haemoglobin and oxygen). § 6 gives a numerical example of a pharmacokinetic system, representing the delivery of a gaseous anaesthetic to a human subject, based on the models developed by WW Mapleson [18–20]. § 7 summarises the paper and suggests further research.

The bond graph modelling in this paper is based on the `BondGraphTools` Python package [21] available at https://pypi.org/project/BondGraphTools/. The code used for the examples in this paper is available at https://github.com/gawthrop/Advection22.

## 2 Hydrochemical transduction

Although different physical and chemical domains have different quantities and units, they share the same energy with units of Joules (J). This fact is used by the bond graph approach to provide a unified approach to *energy transduction* between different energy domains using the transformer **TF** and gyrator **GY** elements.

This section looks at hydrochemical transduction with unidirectional incompressible hydraulic flow; bidirectional flow is considered in § 3. In the incompressible hydraulic domain, the two bond graph covariables are pressure *P* [Pa] and volumetric flow *Q* [m^3^ s^−1^] [22]. Noting that the unit Pa can be rewritten as J m^−3^, the product of the covariables has units J s^−1^. In the chemical domain, the two bond graph covariables are *chemical potential μ* [J mol^−1^] and advective molar flow^1^ *f* [m^3^ s^−1^]. The flow variables are related by:

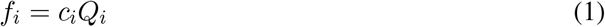

where *c_i_* [mol m^−3^] is the *concentration* of the substance in the liquid.

Figure 1 shows the bond graph of a two domain system. The section labelled *Hydraulic* represents the incompressible hydraulic flow alluded to above. The **R**:**r_i_** component represent an orifice though which the fluid flows; the flow *Q* is typically a non-linear function of the net pressure across the orifice [5, 22]. The section marked *Chemical* represents a single substance being carried by the fluid though the orifice.

**Figure 1:**
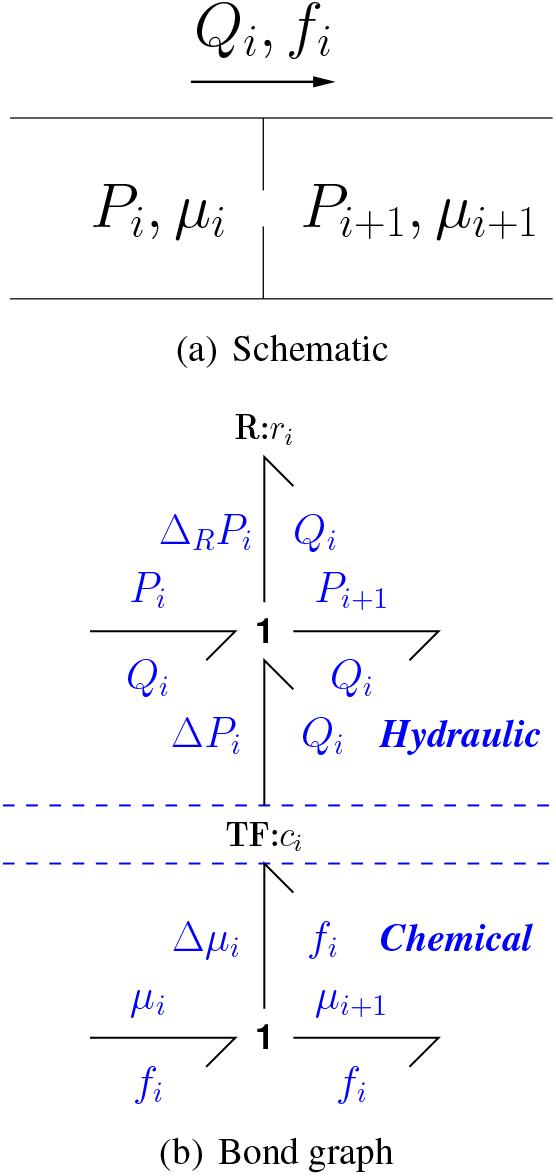
Hydrochemical transduction. (a) Hydraulic flow *Q_i_* through an orifice with pressures *P_i_* and *P*_*i*+1_ and chemical advection with flow *f* and chemical potentials *μ_i_* and *μ*_*i*+1_. (b) The bond graph **R**:**r_i_** component represents the orifice hydraulic resistance and the bond graph modulated transformer component **TF**:**c_i_** transduces energy between the chemical and hydraulic domains with modulus upstream concentration *c_i_*. The bond direction corresponds to energy flow from chemical to hydraulic and *P*_*i*+1_ = *P_i_* + Δ*P_i_* – Δ_*R*_*P_i_* and *μ*_*i*+1_ = *μ_i_* – Δ*μ_i_*. Note that the total pressure drop *P_i_* – *P*_*i*+1_ might not be the same sign as the total chemical potential drop *μ_i_* – *μ*_*i*+1_ due to the pressure drop Δ_*R*_*P_i_* across the hydraulic **R**:**r_i_** component.

The hydraulic and chemical domains are connected by the transformer **TF**:**c_i_** component with modulus *c_i_* which ensures that the chemical flow *f* and hydraulic flow *Q* are related by Equation (1). As the bond graph component transformer **TF**:**c** transmits, but does not dissipate, energy it also follows that:

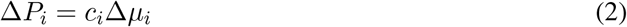

With reference to Figure 1, the energy flow, or power ***P***_*TF*_, transferred by the transformer **TF**:**c** is the product of the flow and effort variables

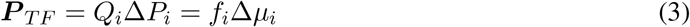

The power dissipated in the **R**:**r_i_** component is:

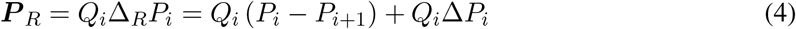

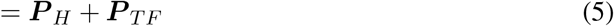

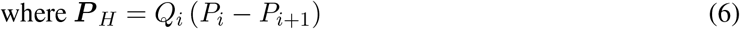

Thus the power dissipated in the **R**:**r_i_** component is the sum of the hydraulic dissipation ***P***_*H*_ and the power ***P***_*TF*_ transmitted from the chemical domain.

## 3 Advection

In this section, it is assumed that, although the hydraulic flow though the orifice strongly affects the chemical flow, the chemical potential has negligible effect on the hydraulic pressure across the orifice. It is therefore possible to approximate the hydraulic dynamics by neglecting the chemical dynamics and, as discussed in this section, the chemical dynamics can be reformulated in terms of a modulated resistance, thus bringing *transport* of a chemical via advection within the same conceptual framework as *transformation* via a chemical reaction.

If the hydraulic flow *Q_i_* is positive, the advective flow *f_i_* is given by Equation (1); if hydraulic flow *Q_i_* is negative, the advective flow *f_i_* is dependent on the concentration *c*_*i*+1_ rather than *c_i_*. Moreover, the chemical potential *μ_i_* is given by [9]:

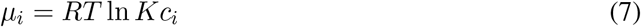

Hence the advective flow *f* is given by

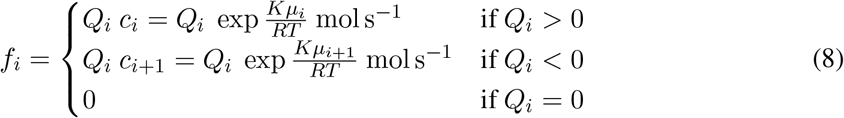

Equation (8) can be modelled as the constitutive relation of a bond graph modulated (by hydraulic flow *Q_i_*) resistive **R** component; to reflect the special properties arising from Equation (8) this is given a special name: the **RA** (advective **R**) component. This component is compared and contrasted with the chemical transformation **Re** component [9] in § 4.

The energy flow, or power ***P***_*RA*_ dissipated by the advective **R** component **RA** is the difference between the product of the flow and effort variables on the two bonds:

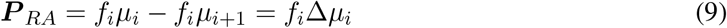

From Equation (3), it follows that ***P***_*RA*_ = *P_TF_* and thus the power dissipation associated with the **RA** component in the representation of Figure 2 corresponds to the power transferred by the **TF**:**c** component, and then dissipated in the **R**:**r_i_** component, in the representation of § 2, Figure 1 and Equation (4). Note that since the transport of chemical is driven by the hydrualic flow, ***P***_*RA*_ may be negative, in contrast to a typical **Re** component which always has non-negative power dissipation.

**Figure 2:**
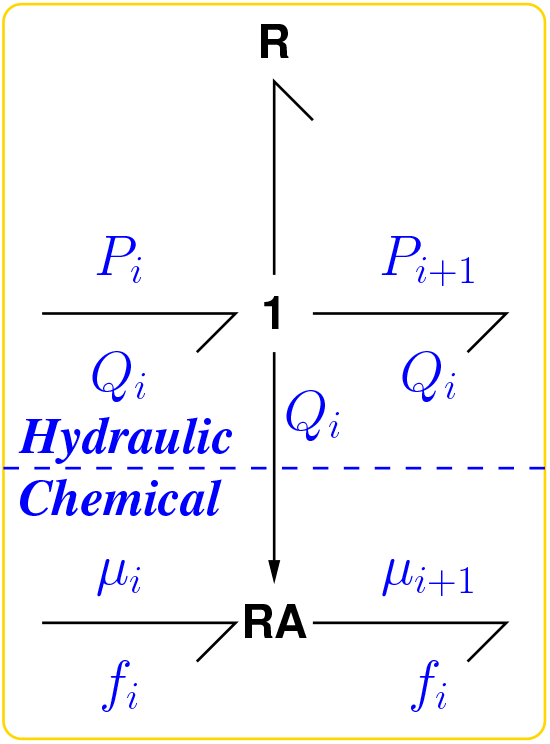
Advection. The transduction of Figure 1 is approximated by a modulated resistive component **RA** (equation (8)) giving one-way interaction from the hydraulic domain to the chemical domain to represent advection of the chemical species by the fluid.

### 3.1 Flow through a pipe

Although the flow of incompressible fluid though an elastic pipe can be modelled by a PDE, it is convenient to use a lumped model instead; this has been examined in the context of blood flow by Safaei et al. [17]. Figure 3(a) shows a lumped model where the pipe is divided into a number of compartments and Figure 3(b) gives the corresponding bond graph. The hydraulic-chemical interaction is one-way, via the inter-compartmental hydraulic flows *Q_i_*; thus the chemical part of Figure 3(b) does not depend on the details of the hydraulic model; only the flows are required. Thus, for example, inertial terms could be added to the hydraulic model by appending bond graph **I** components to the **1** junctions [17, 22].

**Figure 3:**
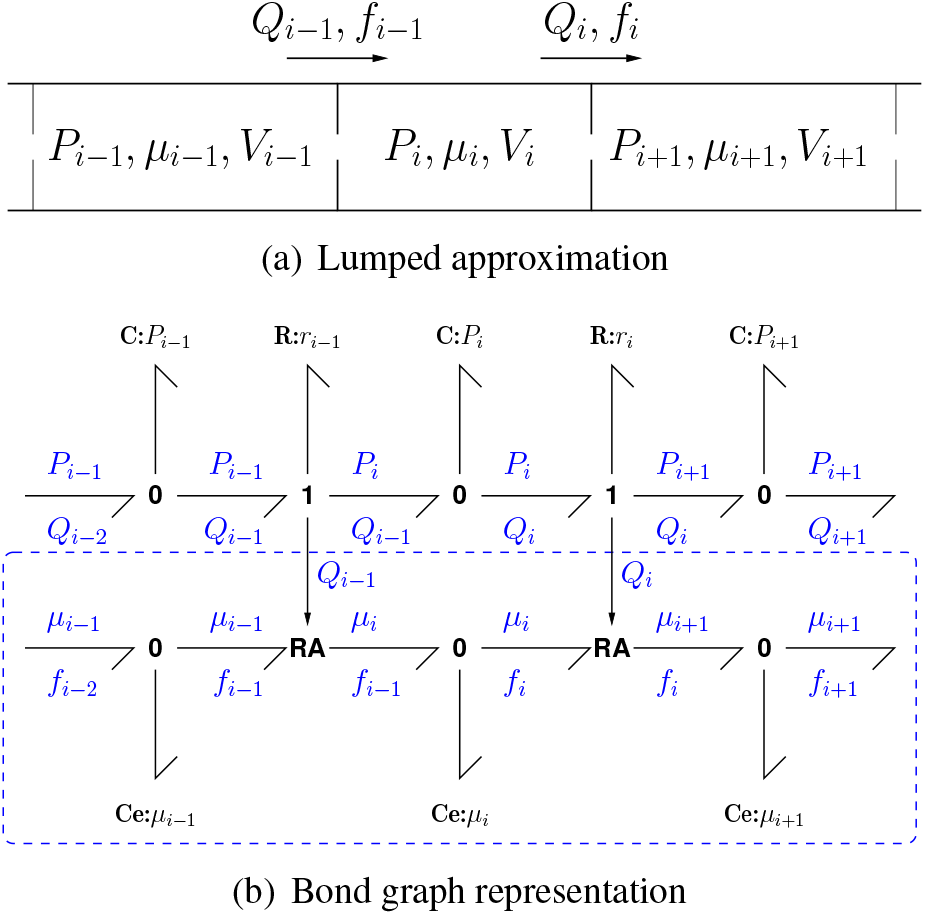
Flow through a pipe. (a) The hydraulics are approximated by a series of compartments of (possibly time-varying) volume *V_i_* containing fluid at pressure *P_i_* and a chemical with chemical potential *μ_i_*. *Q_i_* and *f_i_* are the hydraulic and molar flow rates between compartments *i* and *i* + 1. (b) The bond graph representation of the compartmental model; the advection of the chemical by the fluid is represented by the **RA** components of Figure 2. If the flow rates *Q_i_* are already determined, only the portion within the dashed box is required – this is used as the **Pipe** component in §§ 5& 6.

Advection via a pipe introduces a time-delay into the system dynamics, and the lumped approximation of a pipe introduces an approximate pure time-delay. In particular, if the pipe has *N* compartments, the total pipe volume *V* is:

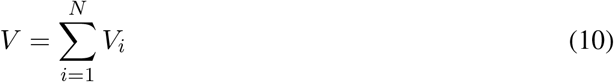

In the case of identical compartments:

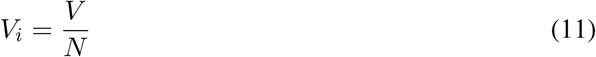

In the case of constant flow *Q*, transfer function analysis shows that

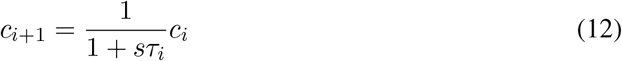

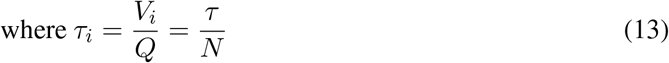

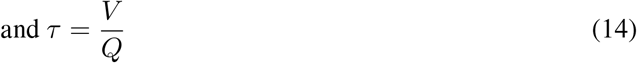

where *s* is the Laplace variable. Hence

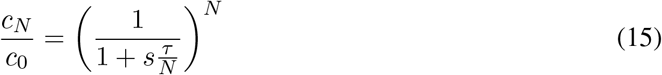

Equation (15) is an approximation to a pure time delay *e*^−*sτ*^; in particular:

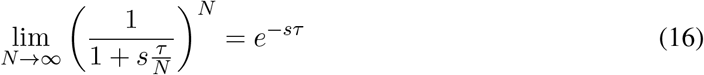

Figure 4 illustrates this approximation.

**Figure 4:**
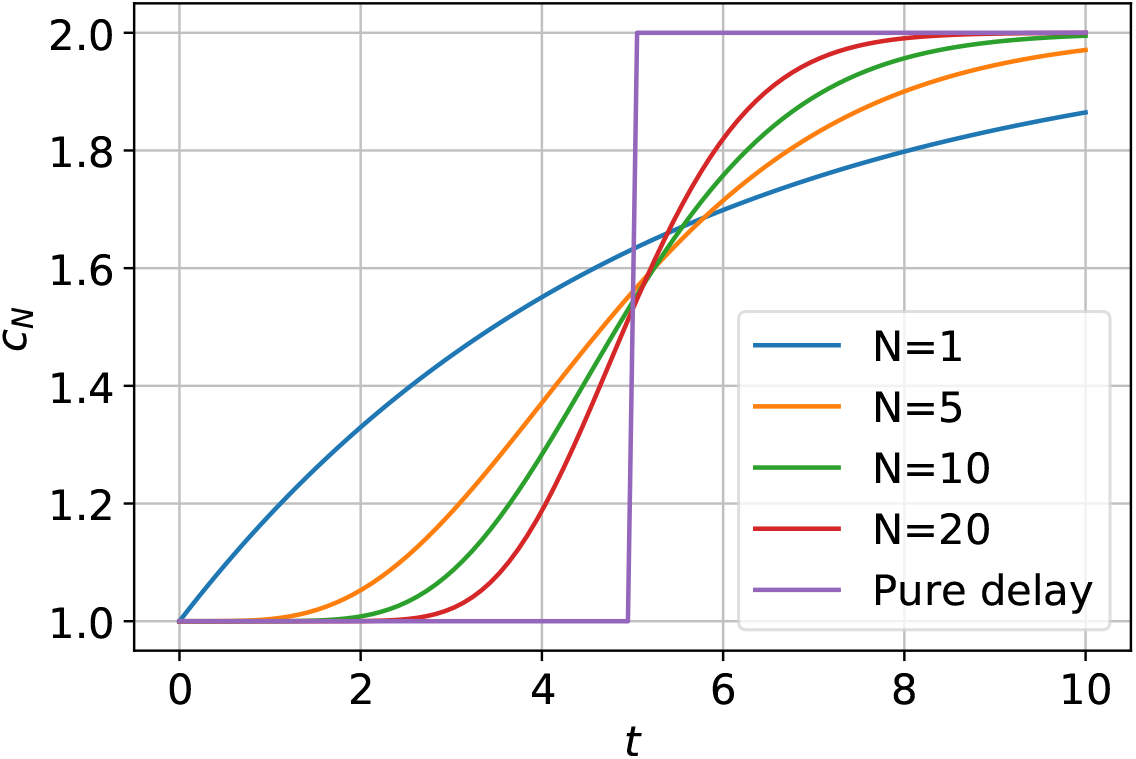
Dynamic response: 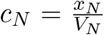. With a constant hydraulic flow-rate *Q* = *Q*_0_, a step change in concentration *c*_0_ at the downstream end of a pipe of volume *V* = 5 modelled with *N* compartments leads to a change in concentration *c_N_* at the upstream end of the pipe. As *N* increases, the response approaches that of a pure time delay of 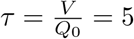.

## 4 Coupled Advection and Transformation

As mentioned in the Introduction, there are two distinct meanings of “molar flow”: the *transport* of a substance from one place to another via advection and the *transformation* of one substance to another via a chemical reaction [10]. As discussed in § 3, advection may be modelled using the **RA** bond graph component together with the capacitive **Ce** components and bonds carrying chemical potential *μ* and molar flow *f*. Moreover, transformation can be modelled by the **Re** bond graph component together with the capacitive **Ce** components and bonds carrying chemical potential *μ* and molar flow *υ* [9]. Hence both concepts can be combined in a single bond graph; this combination is illustrated using a simple example. Further, it is shown how transport via the **RA** component and transformation via the **Re** component can be analysed within a common framework.

For example, Figures 5(a)–(b) correspond to a substance A transformed to substance *C* advected though an orifice with hydraulic flow *Q* into a well-stirred compartment within which the substance C is transformed to substance B. The substance C occurs at two locations so the corresponding **Ce** components are distinguished by denoting them **Ce**:**C**_1_ and **Ce**:**C**_2_; the corresponding amounts of substance (C) in each location are *x*_1_ and *x*_2_ with corresponding concentrations *c*_1_ and *c*_2_.

**Figure 5:**
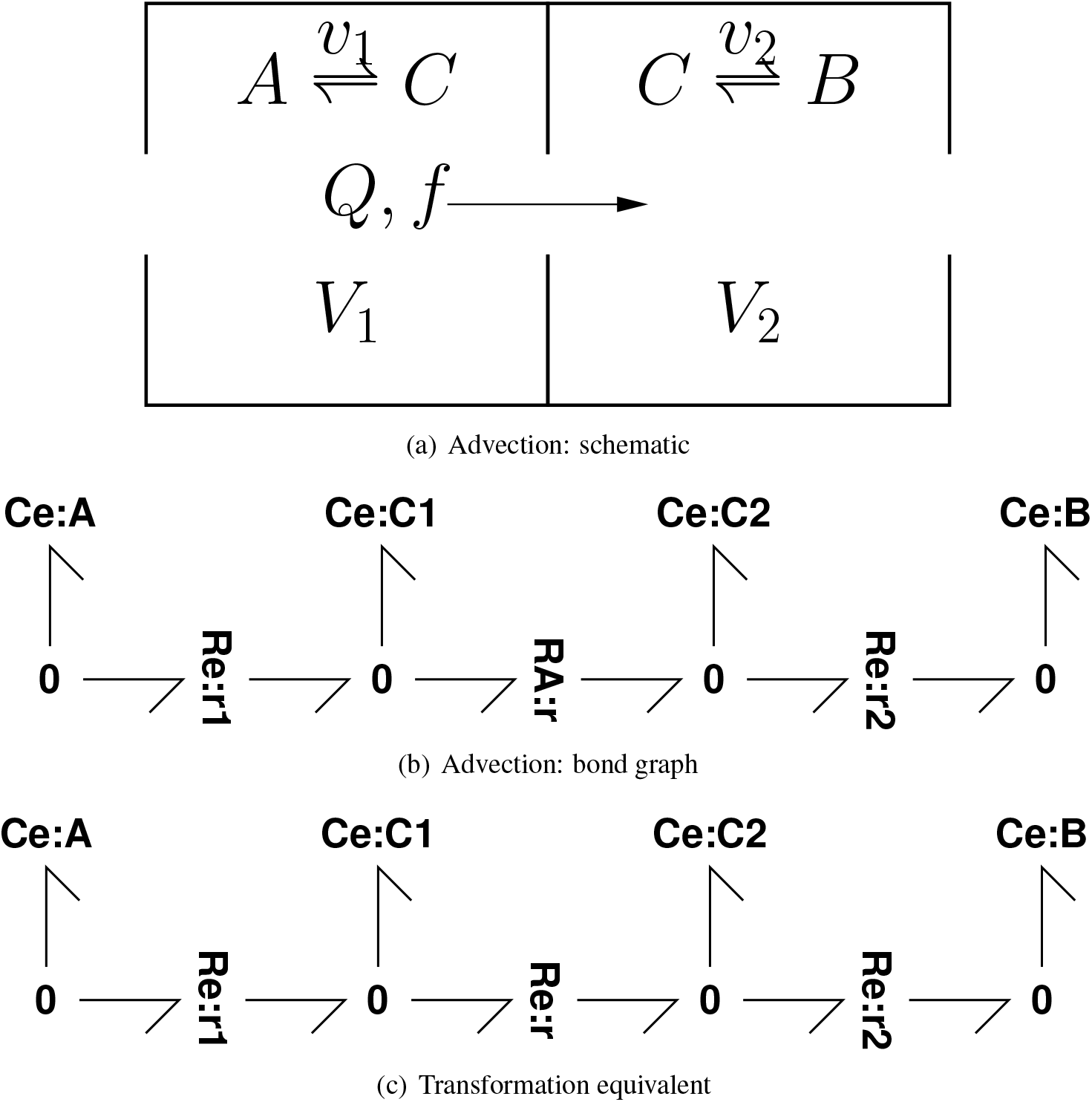
Coupled Advection and Transformation

In the case of positive flow, and in the context of Figure 5(b), Equation (8) becomes

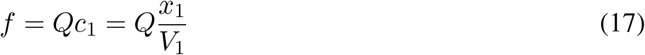

where *f* is the advective flow though **RA**:**r** and *c*_1_ is the concentration corresponding to **Ce**:**C1**. Figure 5(c) is identical to Figure 5(b) except that **RA**:**r** is replaced by **Re**:**r**. In this case, the transformation flow *υ* though **Re**:**r** is [9, Equation 2.2]:

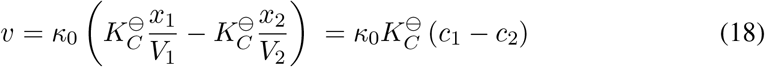

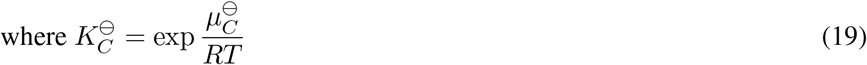

*κ*_0_ is the reaction rate-constant and 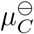 is the standard potential for the species.

Equation (17) can be rewritten as

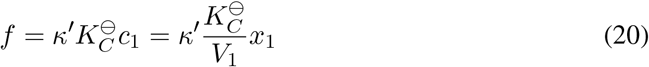

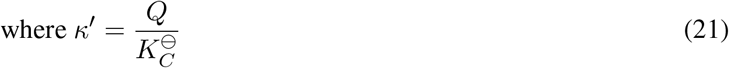

In other words, the **RA** component can be modelled as a **Re** component with flow-dependent rateconstant *κ*′ and a one-way reaction.

Equations (18) and (20) can be combined using the symbol λ where λ = 0 corresponds to Equation (20) for advection and λ = 1 corresponds to Equation (18) for transformation; equations (18) & (22) can be put in a common format as:

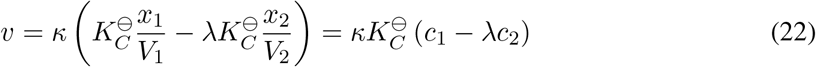

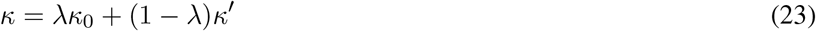

Using this notation, the equations describing both Figures 5(b) and 5(c) become:

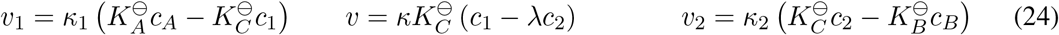

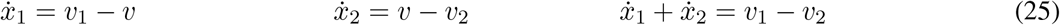

hence

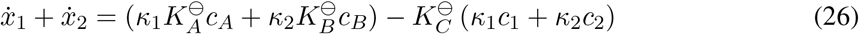

### 4.1 Illustrative example

To illustrate the difference between transportation (λ = 0) and transformation (λ = 1), consider the special case of the system of Figure 5 where the rate constants of the two reactions r_1_ and r_2_ are the same: *κ*_1_ = *κ*_2_.

In the steady-state, the time derivatives are zero and so, using Equation (26):

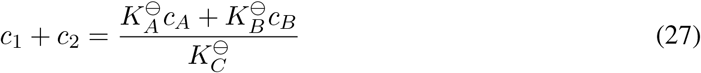

If the flow *Q* is large, *κ* is also large and Equation (24) implies that

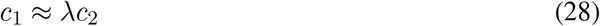

Hence large flow in the steady state implies that:

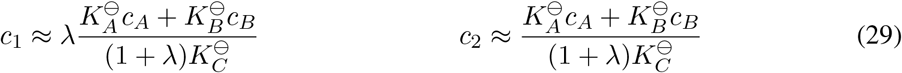

Thus when λ = 0 (advection, Figure 5(b)):

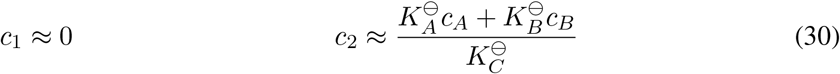

and when λ = 1 (transformation, Figure 5(c)):

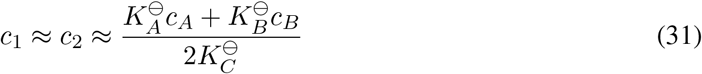

The behaviour for a particular set of values is illustrated by the simulation results of Figure 6. The steady state values for large flow are explained by the analysis leading to Equations (30) and (31).

**Figure 6:**
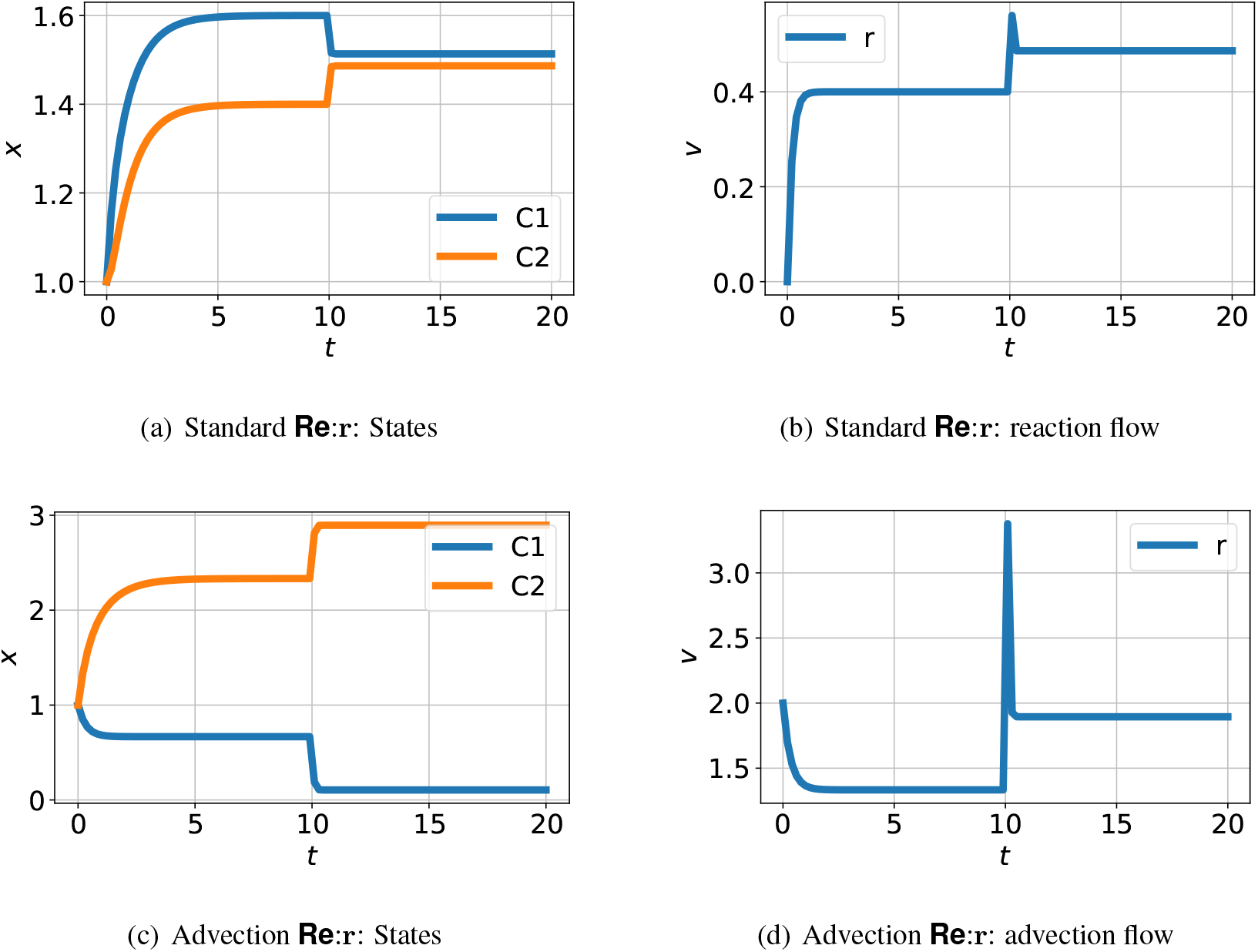
Coupled Advection and Transformation: orifice. (a) amounts *x*_1_ and *x*_2_ when **Ce**:**r** is a reaction component. (b) The flow *υ_r_* corresponding to (a). (c) as (a) but when **Re**:**r** is an advection component. (d) the orifice flow corresponding to (c). The species components **Ce**:**A** and **Ce**:**B** fixed at constant concentations with *x_A_* = 2 and *x_B_* = 1; the species components **Ce**:**C1** and **Ce**:**C2** are free to vary and have unit parameters 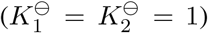. The flow *Q* = 2 when *t* < 10 and *Q* = 10 when *t* ≥ 10. When *Q* = 10, the behaviour corresponds to Equations (30) and (31): (a) 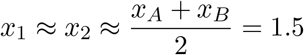; (c) *x*_1_ ≈ 0, *x*_2_ ≈ *x_A_* + *x_B_* = 3.

Figure 7 shows simulation results corresponding to Figure 6 (c) and (d) but with the orifice component replaced by the pipe model of § 3.1. The orifice results are superimposed as a dashed line and show that the steady-state values for the orifice and pipe situations are the same whilst the transient behaviour is different.

**Figure 7:**
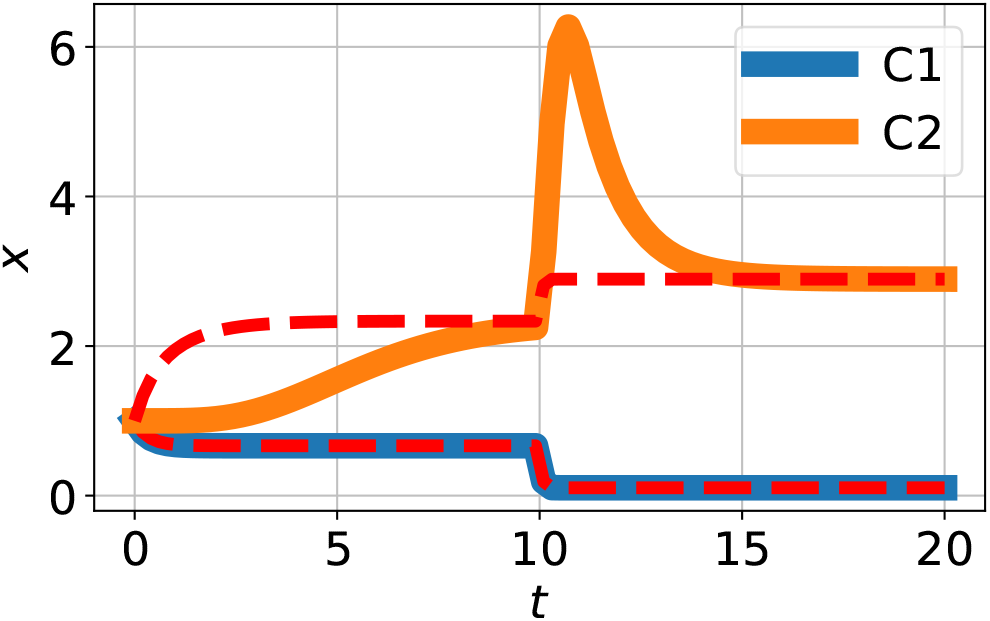
Coupled Advection and Transformation: pipe. The orifice component of Figure 5(b) is replaced by the pipe model of § 3.1 and the orifice results are superimposed as a red dashed line. The behaviour of *x*_1_ is identical to the orifice result of Figure 6(c) as the flow corresponding to the first segment of the pipe is the same as that of the orifice. Due to the pipe, the behaviour of *x*_2_ is delayed with respect to the orifice case but has the same steady-states.

## 5 Circulatory Advection

The cardiovascular systems of humans are closed in the sense that the blood is recirculated round the body: it forms a *circulatory advection* system where the blood advects many substances, notably oxygen (O_2_) and carbon dioxide (CO_2_). There are, in fact a number of circulatory systems including coronary circulation, pulmonary circulation, cerebral circulation and renal circulation. The transport of O_2_ involves the binding and unbinding of O_2_ to haemoglobin which is itself advected by the blood.

Using a simple single circulation model, this paper shows how, in principle, the advection models of § 3.1 and the advection-transformation models of § 4 can be used to build models of a circulatory advection system involving binding and unbinding. More complex systems could be built using the modular properties [6] of bond graphs.

Figure 8 shows a simplified binding/unbinding cycle which has been split into two halves corresponding to two different locations. This section looks at ways in which these two halves can be connected using the advection models developed in § 3. In each case, four advection connections are required to carry both bound C and unbound E enzymes to and from the two locations. Figure 9(a) shows orifice advection connections using four instances of the **RA** component of § 3 whereas Figure 9(b) uses four instances of the **Pipe** component of § 3.1.

**Figure 8:**
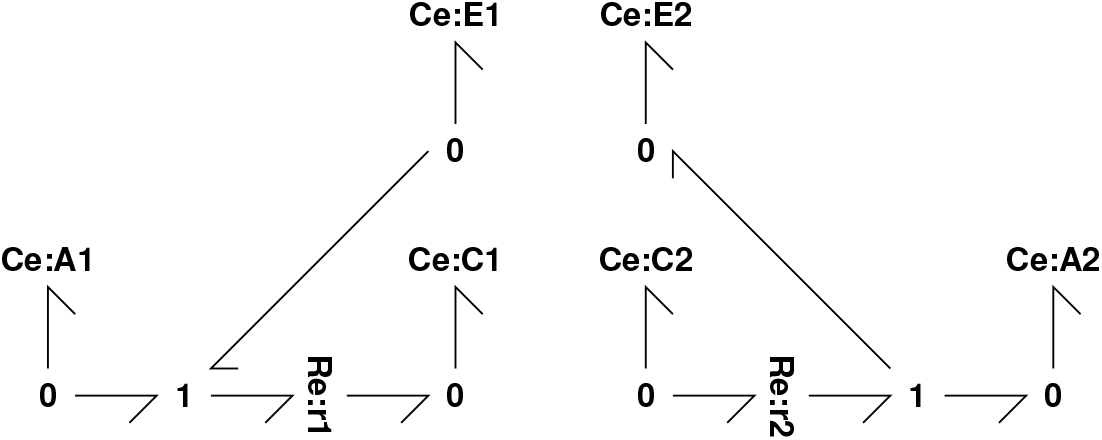
Split binding/unbinding cycle. At location 1, the small molecule A binds to enzyme E to form complex C; at location 2, the complex C unbinds into the small molecule A and enzyme E. C and E are transferred back and forth between the two locations by advection. For example, the small molecule A could represent oxygen and C and E could represent haemoglobin with and without bound oxygen. Location 1 could be the lungs and location 2 an oxygen-consuming organ.

**Figure 9:**
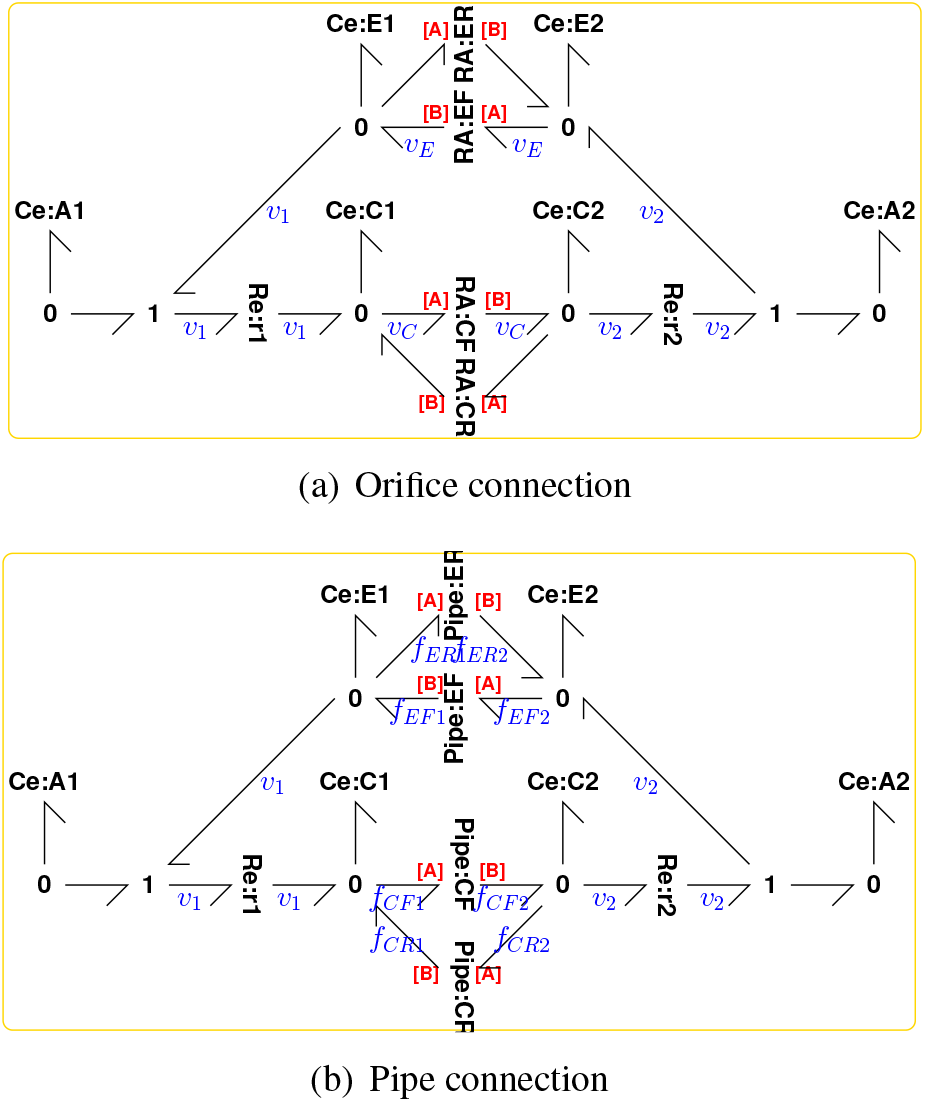
Circulatory Advection. (a) Orifice advection: four instances of the **RA** component of § 3 connect the two halves of the split reaction of Figure 8. **RA**:**CF** corresponds to advection of the bound enzyme complex C from location 1 to 2 and **RA**:**CR** the reverse flow; **RA**:**EF** corresponds to advection of the unbound enzyme E from location 2 to 1 and **RA**:**ER** the reverse flow. (b) as (a) but using the **Pipe** component of Figure 3(b), § 3.1 in place of the **RA** component.

Figure 10 gives some simulation results with unit parameter values except for volume *V* = 5 and (in the case of the pipe connection) number of lumps *N* = 5.

**Figure 10:**
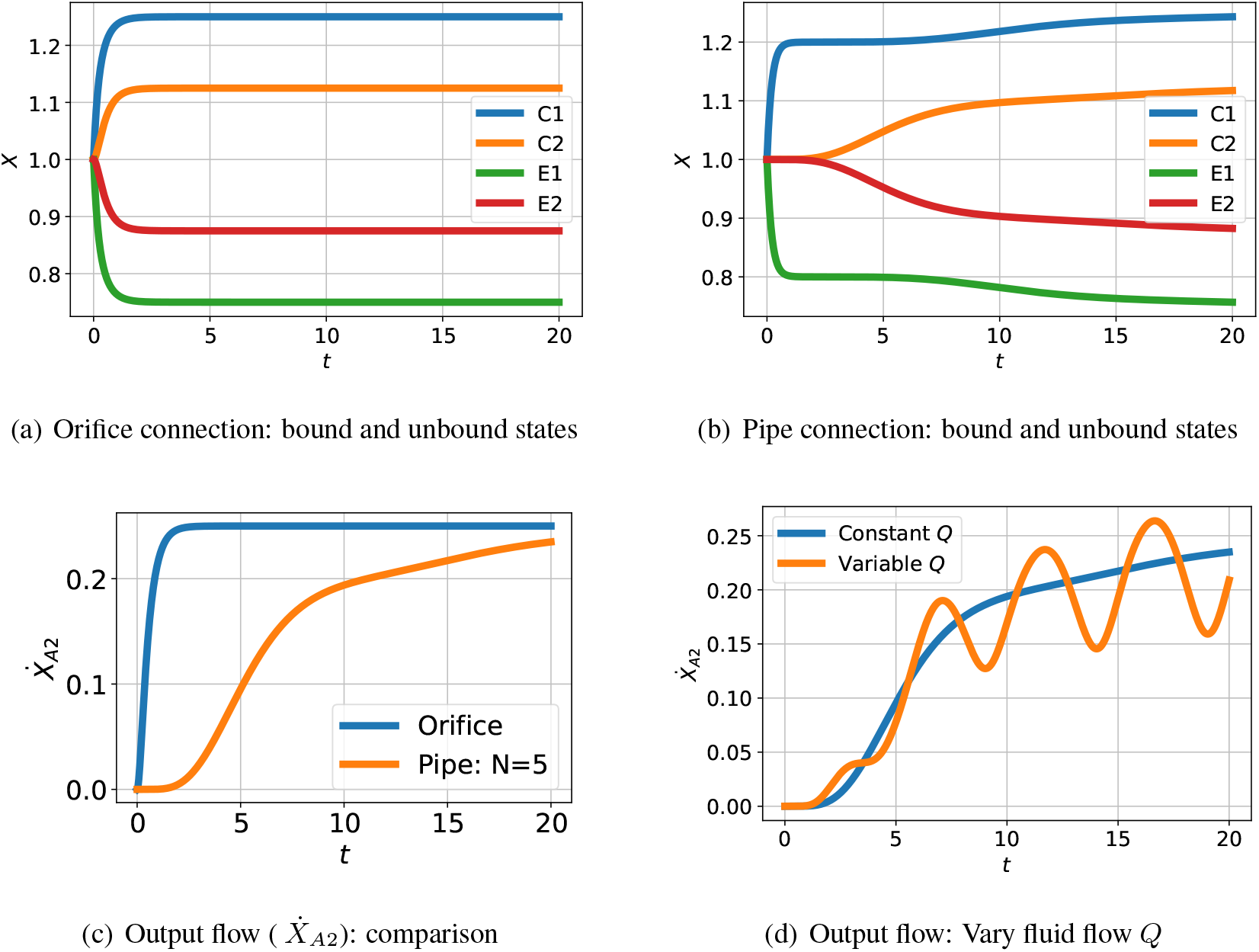
Simulation. For the purposes of illustration, the species A1 and A2 are held at constant concentrations with *x_A_* = 2 and *x_B_* = 1. (a) Orifice connection (Figure 9(a)): Bound *x*_*C*1_&*x*_*C*2_) and unbound (*x*_*E*1_&*x*_*E*2_) states evolving with time *t*. (b) As (a) but with pipe connection (Figure 9(b)) modelled with *N* = 5 lumps; the response is delayed with respect to (a). (c) The flow of substance A (*Ẋ*_*A*2_) into location 2 for cases a) and b); again, the effect of the pipe is to delay the response by about *τ* = 5. (d) The output flow (*Ẋ*_*A*2_) corresponding to (b) with periodically varying fluid flow *Q* = 1 + 0.5 sin 2*πt/T* with period *T* = 5.

## 6 Pharmacokinetics

Pharmacokinetics is the study of drug uptake in living creatures; for example, the anaesthetic gas nitrous oxide (N_2_O) is administered during operations by adding it to inspired air. As it acts on the brain, the dynamics of the transfer of N_2_O from the lungs to the brain via blood circulation is of interest. In his classic papers, Mapleson [18, 19] gave a fully parameterised compartmental model of the uptake of the anaesthetic gas N_2_O in humans outlined in Figure 11. The eight compartments are listed in Figure 11; arterial blood flows from the lungs to the seven organ compartments and returns as venous blood. A simple bond graph model was given by Worship [23] and Gawthrop and Smith [24, Chapter 9]. Rather than model the forking arteries and veins, the blood flow is approximated by a separate arterial flow to, and venous flow from, each compartment; the fraction *δ_p_* of blood passing into each compartment is listed in Table 1. In this section, the bond graph model of [24] is extended to explicitly include the dynamics of the veins and arteries using the approach of § 3.1.

**Figure 11:**
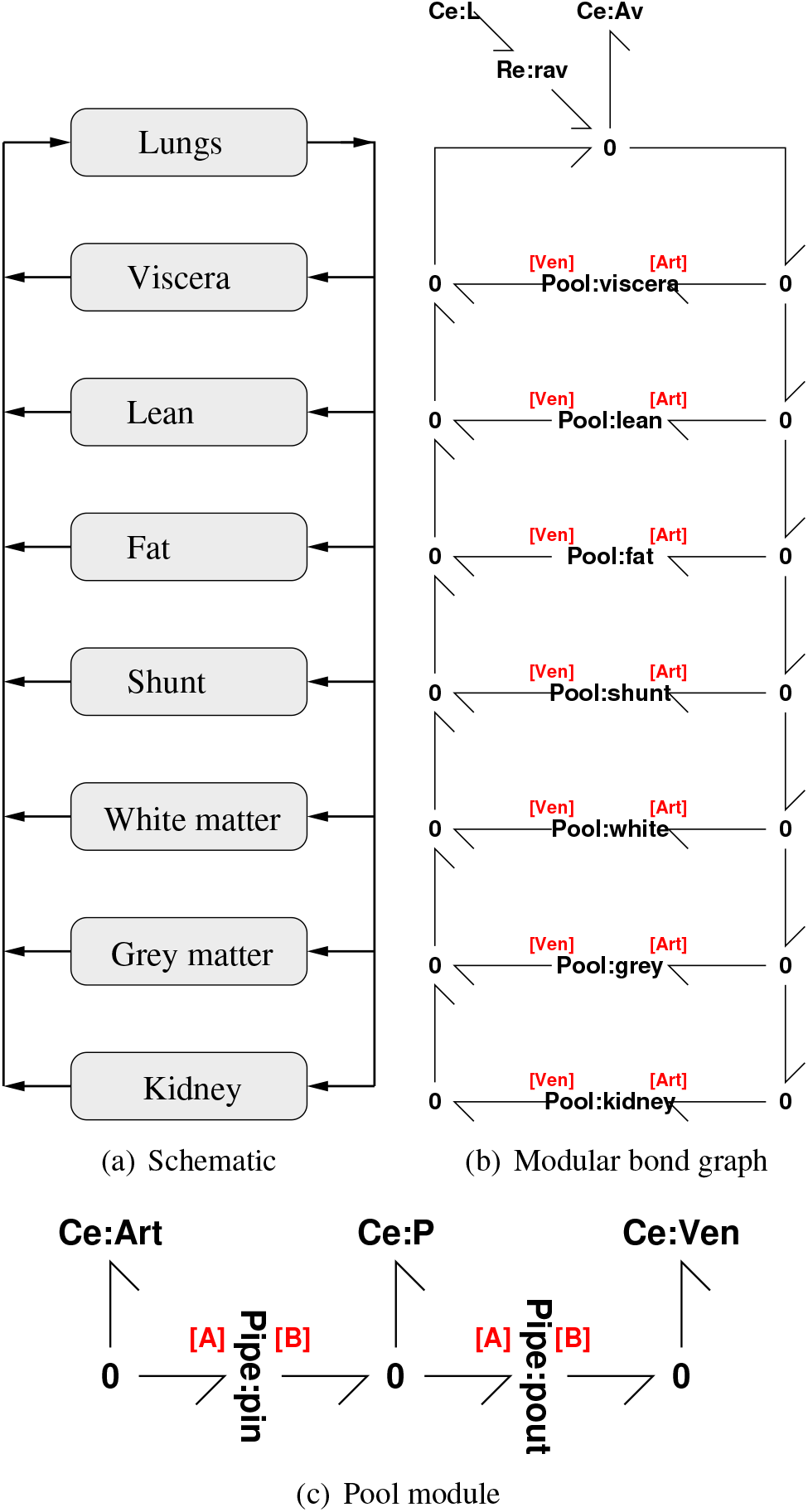
Pharmacokinetics: (a) The schematic diagram shows how the lungs are connected to six compartments representing organs together with a shunt representing blood bypassing organs [18, 19]. (b) A modular bond graph representation of (a). The Pool module appears in (c) and 7 instances appear. [Art] and [Ven] correspond to the two module ports conveying arterial and venous blood respectively. The diffusion of nitrous oxide (N_2_O) from the lung air (modelled by **Ce**:**L**) to the alveolar blood (modelled by **Ce**: **Av**) is modelled by the **Re**:**rav** component. The **0** junctions distribute the arterial flow and combine the venous flows. (c) A modular representation of a pool. The two instances of the pipe component (pin and pout) correspond to arterial flow in and venous flow out.The **Ce**:**P** component models the capacity of the pool for nitrous oxide (N_2_O).

**Table 1:**
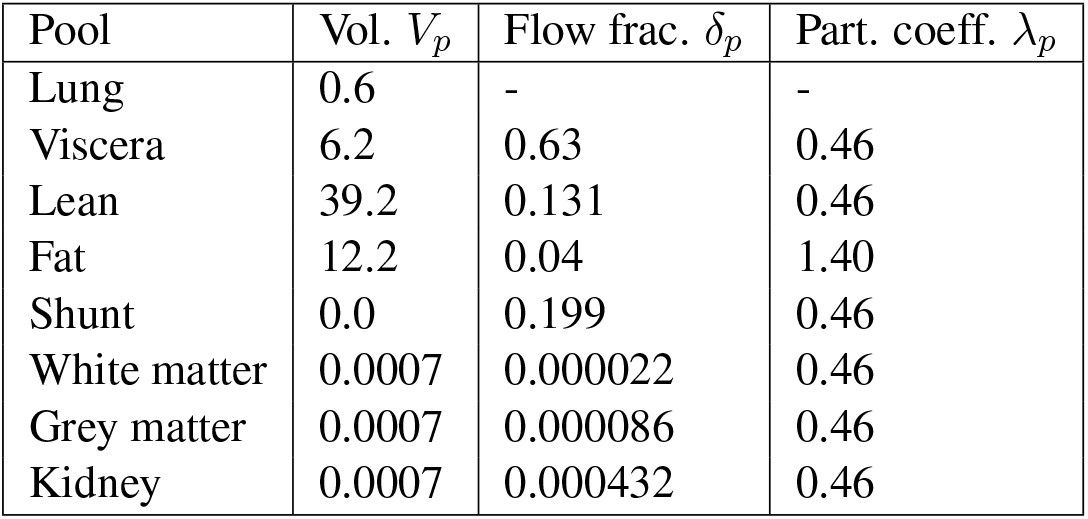
A Detailed Pharmacokinetic Model: data taken from Tables I and II of Mapleson [19]. The volumes of arterial and venous blood are *V_A_* = 1.41 and *V_V_* = 4.01 respectively and the blood flow rate *Q* = 6.481 min^−1^.

For each organ, the compartment **Ce** component has a constant *K_p_* given by

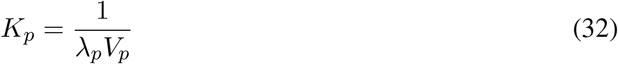

As discussed in § 3.1 and with reference to Table 1 the pipe parameters are:

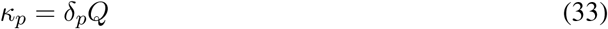

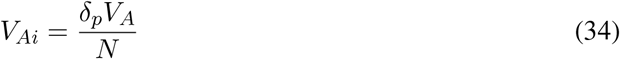

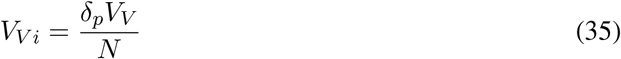

The modular bond graph model of Figure 11 was simulated and the results appear in Figure 12. The results shown in Figure 12(a) correspond closely to those given by Mapleson [19]. The results shown in Figure 12(b) indicate that N_2_O is stored in the fat; the slow release of N_2_O from the fat into the bloodstream can lead to unwanted post-operative anaesthesia.

**Figure 12:**
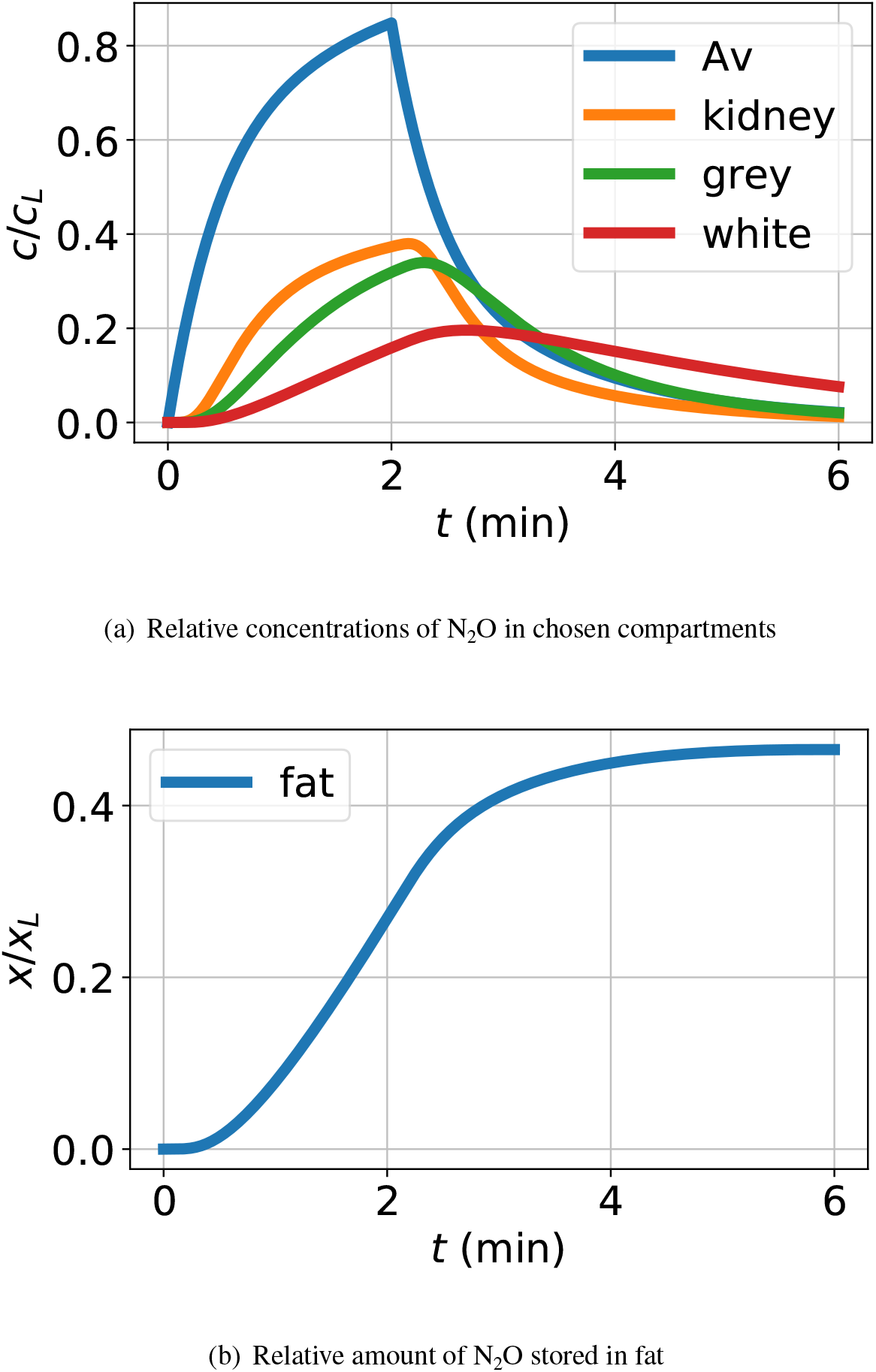
Simulation. Following Mapleson [19], N_2_O is added to inspired air for the first two minutes; for the rest of the simulation pure air is inspired. (a) The concentration of N_2_O, normalised by the initial concentration in the inspired air, is plotted for the alveolar blood and the kidney, brain grey matter and white grey matter compartments. As can be seen, the N_2_O diffuses into and out of the blood via the alveoli and the concentration rises for the first two minutes and then falls as N_2_O is withdrawn. Having passed though the arteries, the arterial N_2_O then drives the individual compartments and is removed by the venous blood. These results are similar to those of Mapleson [19, Figure 2]. The effective time-delay of the arteries is *τ* = *V_A_/Q* ≈ 13 s = 0.22 min; this delay is reflected in the initial response of the three compartmental concentrations. (b) N_02_ accumulates in the fat and takes a long time to decay (beyond the range of this simulation).

## 7 Conclusion

We have shown how advection can be incorporated into bond graph models of physiological systems, providing a physically-consistent framework for linking advective flows with the fluxes associated with chemical reactions. This approach could be used to understand the function of a wide range of organs. Some examples include the transport of oxygen in the blood via haemoglobin; the transport of nutrients and drugs through the liver; and the transport of gases through the lungs. We believe that our approach will allow the incorporation of chemical transformation in these systems as well as transport [17, 25, 26]. Such models could find use in clinical contexts; for example predicting the functional consequences of surgically removing parts of the liver for cancer treatment [27] and understanding the effects of pulmonary obstruction on lung function [28, 29].

Because bond graphs have already been used to model biochemical reactions [9], the models in this paper could be extended to incorporate tissue metabolism as well as advection. This has potential applications in studying the processing of nutrients by the liver and its consumption by tissue [30], as well as in understanding the metabolism and clearance of drugs [31, 32]. The approach could also be used to model complex biochemical kinetics, for example, the cooperative binding/unbinding of oxygen to haemoglobin [33].

A benefit of the approach in this paper is that the building blocks are modular. If a finer level of detail is required, one could replace the simple models of circulation in Figure 11 with more realistic models of vasculature. Furthermore, several organs have a repeating hierarchical structure that can be exploited by a modular approach. For example, the lung is composed of branching pipes that conduct the flow of gas into millions of alveoli [28]. Similarly, the liver is composed of lobes which are in turn made up of several liver lobules [27]. Once bond graph models of fundamental building blocks have been developed, they can be reused in constructing models of more complex anatomical structures

[5]. Such an approach could prove valuable for developing the multiscale models required for the Physiome Project.

To improve the realism of the models presented here, there are some issues that require further investigation:

1. We have assumed that the chemical domain has negligible effect on the hydraulic domain (§ 3). Whilst this is in line with prior approaches [34], the validity of this assumption needs further investigation.
2. This paper considers incompressible flow through rigid pipes; in the case of blood flow the pipes are not rigid, requiring extension to compartments with variable volumes.
3. Hydraulic flow reversal leads to the switching function of Equation (8). This requires further analysis, possibly using switched bond graph methods [35].
4. Species may be carried in cells within the fluid stream – for example, haemoglobin is carried by red blood cells, which may require the modelling of separate compartments.

## Acknowledgements

PJG would like to thank the Faculty of Engineering and Information Technology, University of Melbourne, for its support via a Professorial Fellowship. MP is supported by a Postdoctoral Research Fellowship from the School of Mathematics and Statistics, University of Melbourne.

## Footnotes

1 In this paper, the symbol *υ* is used to represent *transformation*, and *f* to represent *advective*, molar flow; both have the same units: mols^−1^.

## Notes

https://github.com/gawthrop/Advection22

### Competing Interest Statement

The authors have declared no competing interest.

